# Overlapping and unique neural circuits are activated during perceptual decision making and confidence

**DOI:** 10.1101/439463

**Authors:** Jiwon Yeon, Medha Shekhar, Dobromir Rahnev

## Abstract

The period of making a perceptual decision is often followed by a period of confidence generation where one rates the likely accuracy of the initial decision. However, it remains unclear whether the same or different neural circuits are engaged during periods of perceptual decision making and confidence generation. To address this question, we conducted two functional MRI experiments in which we dissociated the periods related to perceptual decision making and confidence report by either separating their respective regressors or asking for confidence ratings only in the second half of the experiment. We found that perceptual decision making and confidence reports gave rise to activations in large and mostly overlapping brain circuits including frontal, parietal, posterior, and cingulate regions with the results being remarkably consistent across the two experiments. Further, the confidence report period activated a number of unique regions, whereas there was no evidence for the decision period activating unique regions not involved in the confidence period. We discuss the possible reasons for this overlap and explore their implications about theories of perceptual decision making and confidence generation.

## Introduction

Perceptual decision making is the process of making a judgment about the identity of a stimulus based on the available sensory information^1^. This process is engaged, for example, each time we judge the color of a traffic signal or the identity of a person down the hallway. Once our perceptual decision is formed, we are able to evaluate the likely accuracy our this decision using ratings of confidence^2,3^. Confidence judgments are often referred to as ‘metacognitive’ because they represent a second-order decision about the accuracy of a first-order decision^4–6^.

Perceptual decision making and confidence are strongly related to each other. In most computational frameworks, they are conceptualized as two separate judgments made on the exact same underlying information^7–12^. The computational similarities suggest that these processes may be supported by similar brain circuits. Support for this notion has come from animal studies demonstrating that the two judgments can be decoded from the same neurons^7,13–16^. Similar findings have been reported in studies with human subjects using electroencephalography (EEG)^17,18^, combined EEG and functional magnetic resonance imaging (fMRI)^19^, or even investigating single neuron activity^20^.

At the same time, many other experiments imply the presence of dissociable neural circuits for perceptual decision making and confidence. Behaviorally, confidence judgments can be dissociated from the accuracy of the perceptual decision in both humans^21–34^ and monkeys^35^. Other studies suggest that confidence judgments but not perceptual decisions are subject to late metacognitive noise^36–43^. Neurally, studies employing transcranial magnetic stimulation (TMS) delivered to the prefrontal cortex have been able to alter subjects’ confidence ratings, while leaving their perceptual decisions unaffected^40,41,44,45^. Similar dissociations between the primary decision and confidence have been observed in studies of memory^46–48^. Such findings have compelled many researchers to hypothesize that perceptual decision making and confidence are based on partially separate neural mechanisms and circuits.

It thus remains unclear whether most of the neural circuits activated during perceptual decision making and confidence are shared or separate. To address this question, here we examined the brain areas activated during periods of perceptual decision making and the confidence report using fMRI. In Experiment 1, we dissociated the decision and confidence periods by separating the regressors for each period. In Experiment 2, we dissociated perceptual decision making and confidence report further by only including confidence judgments in the second half of the experiment. The two experiments produced remarkably similar results showing a large degree of overlap in the brain regions activated during periods of perceptual decision making and confidence reports. Nevertheless, the overlap between the decision and confidence periods was incomplete: both experiments showed several brain areas that were preferentially activated in the confidence period. We explore several possible interpretations of these findings and discuss their implications.

## Methods

### Subjects

Twenty-five subjects completed Experiment 1 (12 females, average age = 21.4 years, range = 18-32 years, compensated $35 for participation). Forty-two subjects completed Experiment 2 but three subjects were excluded due to very low stimulus sensitivity (d’ < 0.2). Hence, 39 subjects were included in the data analyses (23 females, average age = 21.5 years, range = 18-28 years, compensated $50 for participation). Except for one subject, all subjects were righthanded. Subjects had no history of neurological disorders and had normal or corrected-to-normal vision. The study was approved by the Georgia Tech Institutional Review Board. All subjects were screened for MRI safety and provided informed consent. The study’s method and procedure were carried out according to the declaration of Helsinki.

### Stimulus

In both experiments, subjects judged the direction of motion of white dots (density: 2.4/degree^2^; speed: 5°/s) presented in a black circle (Experiment 1: 2° radius; Experiment 2: 3° radius). In Experiment 1, the black circle was positioned either left or right of fixation (its center was 4° from the center of the screen). On each trial, either 4 (low-coherence) or 8% (high-coherence) of the white dots moved coherently and subjects had to determine the direction of motion (either left or right). We used two different coherence levels to ensure that subjects would use both the lower and higher ends of the confidence scales. In Experiment 2, the black circle was positioned at the center of the screen, the coherence level was individually determined for each subject, and the task was to detect whether there was coherent motion (which was always downward) or not. Each dot had a lifetime between three and five frames (refresh rate of the projector: 60 Hz) and the coherent motion was carried by a random subset of dots on each frame. The screen had gray background color. All stimuli were created in MATLAB, using the Psychtoolbox 3^49–51^.

### Task

The task in Experiment 1 was to indicate the direction of motion (left or right) and provide a confidence rating (Figure 1A). Each trial began with a white fixation cross at the center of the screen. The fixation cross was presented randomly for two or three seconds. Following the fixation cross, we presented a cue indicating the likely side of the screen where the stimulus would be presented. The cue was invalid on 10% of all trials. These invalid-cue trials were considered to be catch trials and were analyzed separately. The duration of the cue was randomly chosen to be .5, 1, 2, or 4 seconds. The subjects were then presented with the moving dots stimulus and asked to judge the direction of motion. The stimulus presentation lasted until a response was selected via a button press. Subjects gave their responses using the index and middle fingers to indicate left and right direction, respectively. For the 90% of all trials that contained a valid cue, after subjects provided their response, a prompt to report confidence was presented on 55% of all trials. For the remaining 35% of all trials, the cue was valid but the confidence prompt was not presented, and subjects did not make a confidence response. Confidence was never given on catch trials. The prompt indicated the confidence scale on which confidence should be rated on each trial. We randomly alternated between 2-point, 3-point, and 4-point scales. The scale to be used was signaled by presenting the number 2, 3, or 4 as the confidence prompt. The lowest level of confidence was always indicated by pressing a button with the index finger. There was no time pressure for either the decision or confidence responses.

**Figure 1.**
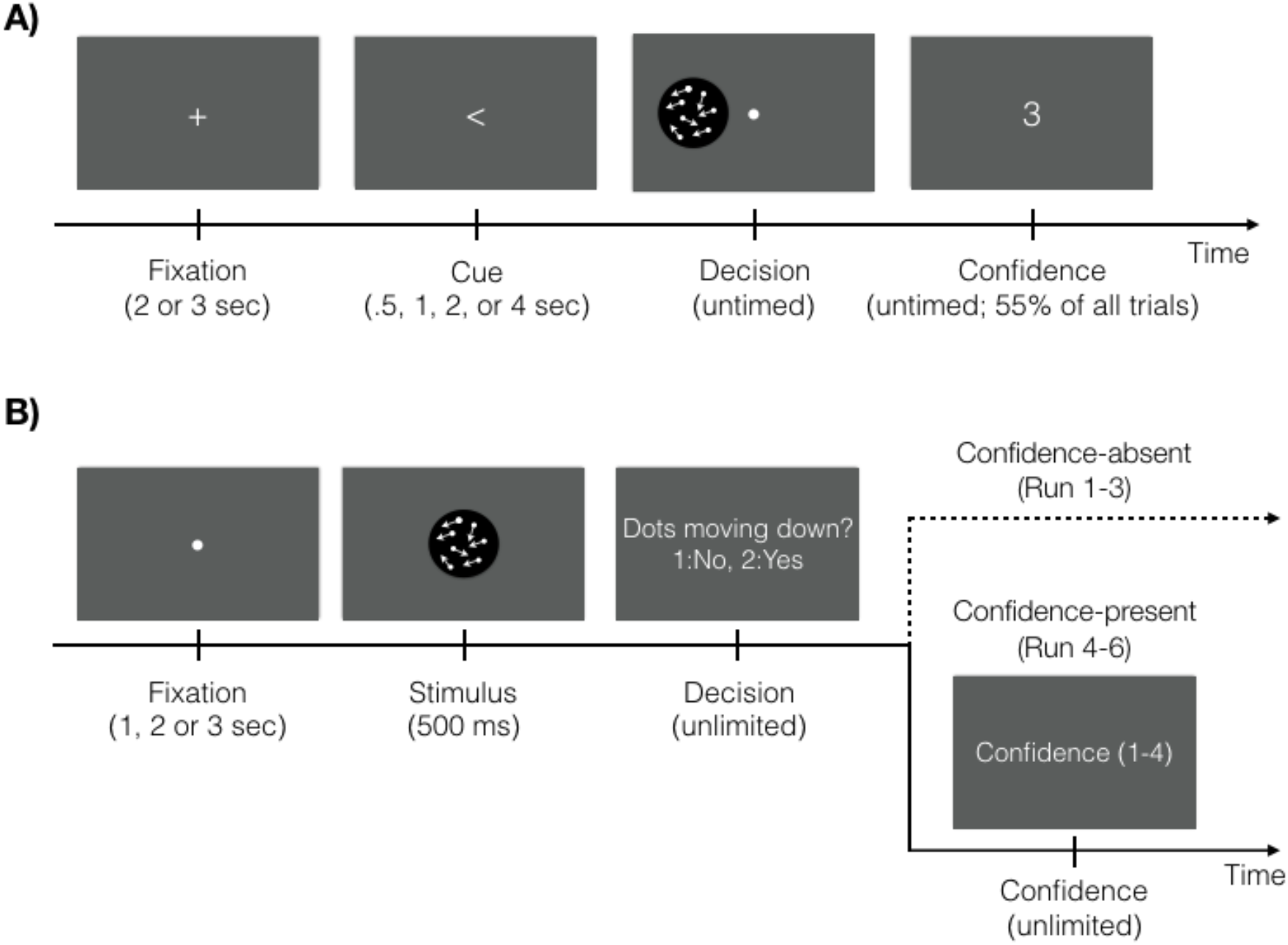
Tasks. (A) Each trial started with a white fixation cross. A cue then indicated the likely location of the following stimulus (either left or right of fixation). In the decision period, the subjects were asked to judge the direction of coherently moving dots. A confidence prompt was presented on 55% of all trials. Three rating scales (2-, 3-, and 4-point scales) were used and the scale was indicated by the number presented on the screen (3-point scale in this figure). (B) Each trial started with a white fixation mark, followed by a moving-dots stimulus presented for 500 ms. Subjects indicated whether the dots moved coherently in downward direction or not. In the first half of the experiment (Runs 1-3), subjects were not asked to evaluate their confidence level (Confidence-absent condition). In the second half of the experiment (Runs 4-6), subjects had to rate their confidence level on a 4-point scale (Confidence-present condition). Subjects were first instructed about giving confidence after completing the Confidence-absent condition.

The task in Experiment 2 was to indicate whether a moving-dots stimulus had an overall coherent motion (always in downward direction) or not (Figure 1B). Each trial began with a fixation mark presented randomly for 1, 2, or 3 seconds. A moving-dots stimulus followed and was presented for 500 ms. After the stimulus offset, subjects indicated whether they detected downward motion by using their right index and middle fingers to respond ‘No’ and ‘Yes,’ respectively. The experiment was separated in two halves. In the first half (Runs 1-3), subjects simply performed the task and were never told anything about evaluating their confidence level; we call this the Confidence-absent condition. In the second half of the experiment (Runs 4-6), subjects made their perceptual decision and immediately after were asked to indicate their confidence level; we call this the Confidence-present condition. Importantly, subjects were first informed about the confidence judgments during a short practice run after the end of Run 3 (that is, after completing the Confidence-absent condition). Confidence was always given on a 4-point scale, where the index and the little fingers indicated the lowest and the highest confidence level, respectively. Subjects had unlimited time for both the perceptual decision and confidence judgments.

### Procedure

Both experiments started with a short training outside the scanner. During the training, the subjects received instructions on how to perform the task and completed 40 example trials. For Experiment 2, we then determined subjects’ threshold coherence level using a 2-down-1-up staircase procedure and presented 60 additional practice trials with the target coherence level. Following the training in both experiments, subjects were positioned in the scanner where we first collected a structural scan. While collecting the structural scan, subjects were given additional practice trials (excepts for two subjects in Experiment 1 due to technical issues). In Experiment 2, we measured subjects’ coherence threshold again using the same 2-down-1-up staircase procedure and then used the threshold in an additional practice block. Based on the performance in this block, the coherence level was adjusted manually for some subjects (mean coherence level = 17%, SD = 5%; mean accuracy = 74.1%, SD = 8%). All responses were given with the right hand via an MRI-compatible button box.

Experiment 1 had four runs, each consisting of four 20-trial blocks (for a total of 320 trials). Blocks were separated by 15-second breaks that were indicated by a black fixation cross at the center of the screen. Subjects were given untimed rest periods between runs. Two subjects completed only three runs and one subject completed three runs and a single block from the fourth run. The remaining 22 subjects completed the full four runs.

Experiment 2 consisted of six runs (Run 1-3: Confidence-absent condition; Run 4-6: Confidence-present condition), with each run containing nine 16-trial blocks. The experiment consisted of three types of blocks – a random motion block (randomly moving dots were more frequent), a downward motion block (downward moving dots were more frequent), and a neutral block (both stimuli were presented equally often). The identity of each block was indicated by a cue that appeared before the block. Within a run, the order of the blocks was pseudorandomized such that the three types of blocks always appeared together in the same random order. However, since the current study is not focused on the effects of cuing, we only included the neutral blocks in our analyses (288 trials in total). Subjects were given untimed rest period between runs. Two subjects completed only five runs and the rest 37 subjects completed all six runs.

### Separating the regressors for perceptual decision making and confidence in Experiment 1

The design of Experiment 1 was optimized to separate the regressors for the cue, decision, and confidence periods. The separation between regressors was achieved via several design features. First, we presented the cue for .5, 1, 2, or 4 seconds (duration was randomly chosen on each trial), thus separating the regressor for the cue period from the regressors for the decision and confidence periods. Second, we used moving dots as stimuli and placed them in the periphery in order to slow down the perceptual decision-making process and ensure that the decision period was as long as possible. Third, we varied the confidence scale across trials, which slowed down the confidence report process thus making that period longer too. Because the contribution of each event is calculated after a convolution with the sluggish hemodynamic response function, longer events are easier to separate from shorter ones such that two short events in succession would not produce differentiable neural activations but two long events would^52–55^. Therefore, elongating the perceptual decision and confidence periods helps separate the regressors associated with these two events. Finally, we only asked for confidence responses on just over half of all trials thus also helping separate the regressors for decision and confidence periods. Overall, these design characteristics resulted in low correlations between the cue, decision, and confidence period regressors (average cue-decision correlation: r = -.15, SD = .064; average cue-confidence correlation: r = -.215, SD = .051; average decision-confidence correlation: r = .16, SD = .105; Supplementary Figures 1 and 2).

However, our attentional manipulation (i.e., the spatial cue indicating the likely location of the stimulus) did not appear to be effective. First, we did not find differences in either accuracy (t(24) = .60, *p* = .56) or reaction time (t(24) = -.17, *p* = .87) between valid and invalid cues. Second, the fMRI activations associated with the cue period did not reveal any of the known brain areas related to spatial attention (e.g., frontal eye fields and intraparietal sulcus). Instead, the cue period appeared to elicit activity in the default mode network (Supplementary Figure 3). These results suggest that subjects did not engage spatial attention during the cue period, and therefore this period could not be used to reveal attention-related processes. We suspect that the reason why subjects employed little to no spatial attention during the cue period is that stimulus presentation was already quite long (average = 1.58 seconds) and hence deploying attention in advance was not beneficial for performance. Because of these considerations, we focused our analyses on the overlapping and unique brain activations produced by the decision and confidence periods.

### fMRI acquisition and preprocessing

The MRI data were collected on 3T MRI systems (Experiment 1: Trio MRI system, Experiment 2: Prisma-Fit MRI system; Siemens) using 12-channel (Experiment 1) and 32-channel (Experiment 2) head coils. Anatomical images were acquired using T1-weighted sequences (Experiment 1: MPRAGE sequence, FoV = 256 mm, TR = 2530 ms, TE = 1.74 ms, 176 slices, flip angle = 7°, voxel size = 1.0 × 1.0 × 1.0 mm^3^; Experiment 2: MEMPRAGE sequence, FoV = 256 mm; TR = 2530 ms; TE = 1.69 ms; 176 slices; flip angle = 7°; voxel size = 1.0 × 1.0 × 1.0 mm^3^). Functional images were acquired using T2*-weighted gradient echo-planar imaging sequences (Experiment 1: FoV = 220 mm, TR = 1780 ms, TE = 24 ms, 37 descending slices, flip angle = 70°, voxel size = 3.0 × 3.0 × 3.5 mm^3^; Experiment 2: FoV = 220 mm; TR = 1200 ms; TE = 30 ms; 51 slices; flip angle = 65°; voxel size = 2.5 × 2.5 × 2.5 mm^3^).

We used SPM12 (Wellcome Department of Imaging Neuroscience, London, UK) to analyze the MRI data. The first two volumes of each run were removed to allow for scanner equilibration. Functional images were first converted from DICOM to NIFTI and then preprocessed with the following steps: de-spiking, slice-timing correction, realignment, segmentation, coregistration, and normalization. The functional images were smoothened with 6 mm full-width-half-maximum (FWHM) Gaussian kernels.

### Analyses

Experiment 1 was initially designed to investigate the shared and unique brain activations produced by the processes of attention, decision, and confidence^40,56^. However, since the attentional manipulation appeared not to be effective (see above), we only focused on analyzing the brain activity elicited by the decision and confidence periods. Experiment 2 was designed with the dual purpose of investigating expectation-related brain activity in detection tasks (these analyses will be reported in a separate publication) and better separating the decision and confidence periods by only introducing confidence in the second part of the experiment. We did not preregister either experiment and hence all analyses should be considered exploratory.

For all contrast tests across the two experiments, the results of individual subjects were submitted to a group-level t-test. Except for the Supplementary Figure 3, all statistical tests were based on *p* < .05 corrected for false discovery rate (FDR) for each individual voxel. We only reported activations of contiguous clusters of at least 150 voxels. All t-values in the Supplementary Tables reflect the peak voxel within the activated clusters.

#### Experiment 1

In a first set of analyses for Experiment 1, we constructed a general linear model (GLM) with 17 regressors for each run. The first six regressors modeled the blood-oxygen level-dependent (BOLD) responses related to the fixation period (spanning the period from onset to offset of the fixation mark), cue period, decision period, catch trials decision period (the same decision period as in the “decision period” regressor but only in catch trials), confidence period, and rest periods between blocks (spanning the period of rest in-between blocks). We modeled catch trials separately since they may invoke additional processes unrelated to perceptual decision making. This and subsequent analyses constitute rapid event-related designs which lead to acceptable levels of separation between regressors in the presence of randomized interstimulus intervals^54,58,59^. In addition, we included six regressors related to head movement (three translation and three rotation regressors), four tissue regressors (white matter, cerebrospinal fluid and bone, soft tissues, and air and background), and a constant term.

In order to determine the brain regions involved in perceptual decision making and confidence report, we compared the decision and the confidence periods to the cue period using the contrasts Decision > Cue and Confidence > Cue. The cue period was chosen as an active control baseline, but the results were very similar if the fixation period was used as a baseline instead. In addition, in order to find brain regions that were preferentially active for perceptual decision making or confidence, we directly compared the decision and the confidence periods (Decision > Confidence and Confidence > Decision).

We further examined whether the results obtained from the analyses above depended on the coherence level or the confidence rating scales. To do so, in a separate GLM analysis, we created separate regressors for the decision period that corresponded to the low- and high-coherence stimuli while keeping all the other regressors the same as in the GLM for the first set of analyses (18 regressors in total). In a separate GLM analysis, we created separate regressors for the confidence period that corresponded to the three different rating scales presented (19 regressors in total). We then ran the same analyses as before for the decision period associated with each coherence level and for the confidence period associated with each confidence scale.

In another control analysis, we separated the decision period regressor into two based on whether a trial contained a confidence report (Decision_with_conf_) or not (Decision_no_conf_). We created a separate GLM analysis, in which we included these two decision-related regressors instead of the single decision regressor as in our first GLM analysis. The other regressors remained the same. We then confirmed that the results remain the same if all of the analyses were performed using the Decision_no_conf_ regressor instead of the original Decision regressor.

The design of Experiment 1 was optimized for separating decision- and confidence-related fMRI regressors. This design did not allow us to compute metacognitive performance or other confidence-related variables because of the inclusion of three different scales for the confidence response (and the corresponding small number of trials related to each scale). Nevertheless, for the purposes of providing summary statistics of subjects’ performance, we normalized each confidence rating such that a confidence rating of *k* given on an n-point scale was transformed to 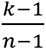. This transformation ensured that the ratings for all confidence scales were in the [0,1] interval. We then averaged the transformed confidence responses to report summary statistics.

#### Experiment 2

We generated two different GLMs in Experiment 2 for the Confidence-absent (Runs 1-3) and Confidence-present (Runs 4-6) conditions. For the Confidence-absent condition, we included regressors related to the fixation period, decision period, and between-block periods. For the Confidence-present condition, we included regressors of the fixation, decision, confidence, and between-block periods. In addition, we included six head-motion related regressors (three translation and three rotation regressors), four tissue regressors (white matter, cerebrospinal fluid and bone, soft tissues, and air and background), and a constant term for all sessions.

We first compared the brain responses related to the decision periods in Confidence-absent (Decision_conf-abs_) and Confidence-present (Decision_conf-pres_) conditions to the fixation period (Decision_conf-abs_ >Fixation and Decision_conf-pres_ > Fixation). Also, to check whether we would observe similar activation patterns for the confidence period, we contrasted the regressors related to the confidence period in the Confidence-present condition to the fixation period (Confidence > Fixation). Furthermore, we examined the unique neural responses related to each decision and confidence periods. Similar to the contrast tests in Experiment 1, we compared the decision period of the two conditions separately with the confidence period from the Confidence-present condition (Decision_conf-abs_ > Confidence, Decision_conf-pres_ > Confidence, Confidence > Decision_conf-abs_, and Confidence > Decision_conf-pres_).

### Data and code

All data and codes for the behavioral analyses are freely available at https://osf.io/pn283 and have also been uploaded to the Confidence Database^60^. In addition, unthresholded fMRI maps have been are uploaded in NeuroVault^61^ and can be accessed at https://neurovault.org/collections/9004/.

## Results

We investigated whether the periods of perceptual decision making and confidence generation activate the same or different brain regions using fMRI. In Experiment 1, we separated the regressors for the decision and confidence periods in an attempt to determine their unique contributions to BOLD activity. In Experiment 2, subjects only made perceptual decisions in the first half of the experiment, but provided both perceptual and confidence judgments in the second half. This design allowed us to ensure that no explicit confidence processes took part during the period of decision making in the first half of Experiment 2. We then compared the brain activations related to perceptual decision making and confidence generation in both experiments.

### Experiment 1

#### Behavioral results

Average task accuracy was 67.6% correct (SD = 9.91), while average response time was 1.58 seconds (SD = .418). Performance was higher for high-coherence (accuracy = 69.5%, SD = 12.1) than for low-coherence stimuli (accuracy = 65.6%, SD = 9.39; paired t-test: t(24) = 2.28, *p* = .032). Similarly, reaction time was faster for the high-coherence task (RT = 1.54 sec, SD = .399) compared to the low-coherence task (RT = 1.63 sec, SD = .444; paired t-test: t(24) = -3.81, *p* = 8.47 x 10^-4^). To compare the confidence ratings across the three different scales (we interleaved 2-, 3-, and 4-point scales), we mapped each scale on the [0, 1] interval such that the lowest confidence rating was always changed to 0 and the highest confidence rating was always changed to 1 (see Methods). The average transformed confidence value across all rating scales was .649 (SD = .191). Subjects were more confident on correct trials (mean confidence = .672, SD = .191) than on incorrect trials (mean confidence = .592, SD = .193; paired t-test: t(24) = 5.217, *p* = 2.40 x 10^-5^). These results suggest that the subjects were able to perform the task as intended and provide appropriate confidence ratings.

#### Shared activations between perceptual decision making and confidence generation

To explore the possible activation overlap elicited by periods of perceptual decision making and confidence generation, we first examined the activations for the decision and confidence periods separately. To do so, we compared the periods of decision and confidence deliberation with the pre-stimulus period when the spatial cue was presented. Our design was successful in decorrelating the cue regressor from both the decision regressor (average r = -.15, SD = .064) and the confidence regressor (average r = -.215, SD = .051) thus allowing us to use the cue period as an active control baseline.

We found that the decision and confidence periods activated a similar set of fronto-parieto-posterior brain regions (Figure 2A,B). In order to examine the amount of overlap, we created a map of the intersection of the decision and confidence period activations (Figure 2C). This map showed extensive bilateral activations in a number of areas including the anterior prefrontal cortex (aPFC), dorsolateral prefrontal cortex (dlPFC), frontal eye fields (FEF), intraparietal sulcus (IPS), and right superior temporal gyrus (STG). In addition, strong bilateral activations were present in the dorsal anterior cingulate cortex (dACC), dorsomedial prefrontal cortex (dmPFC), insula, precuneus, and occipitotemporal cortex (OTC) which includes the motion complex area (MT+). We also observed bilateral somatosensory, motor, and pre-supplementary motor cortex activity, which were likely related to the fact that both the decision and confidence periods featured a button press. Coordinates of peak activity for all regions and corresponding *t*-values are shown in Supplementary Table 1.

**Figure 2.**
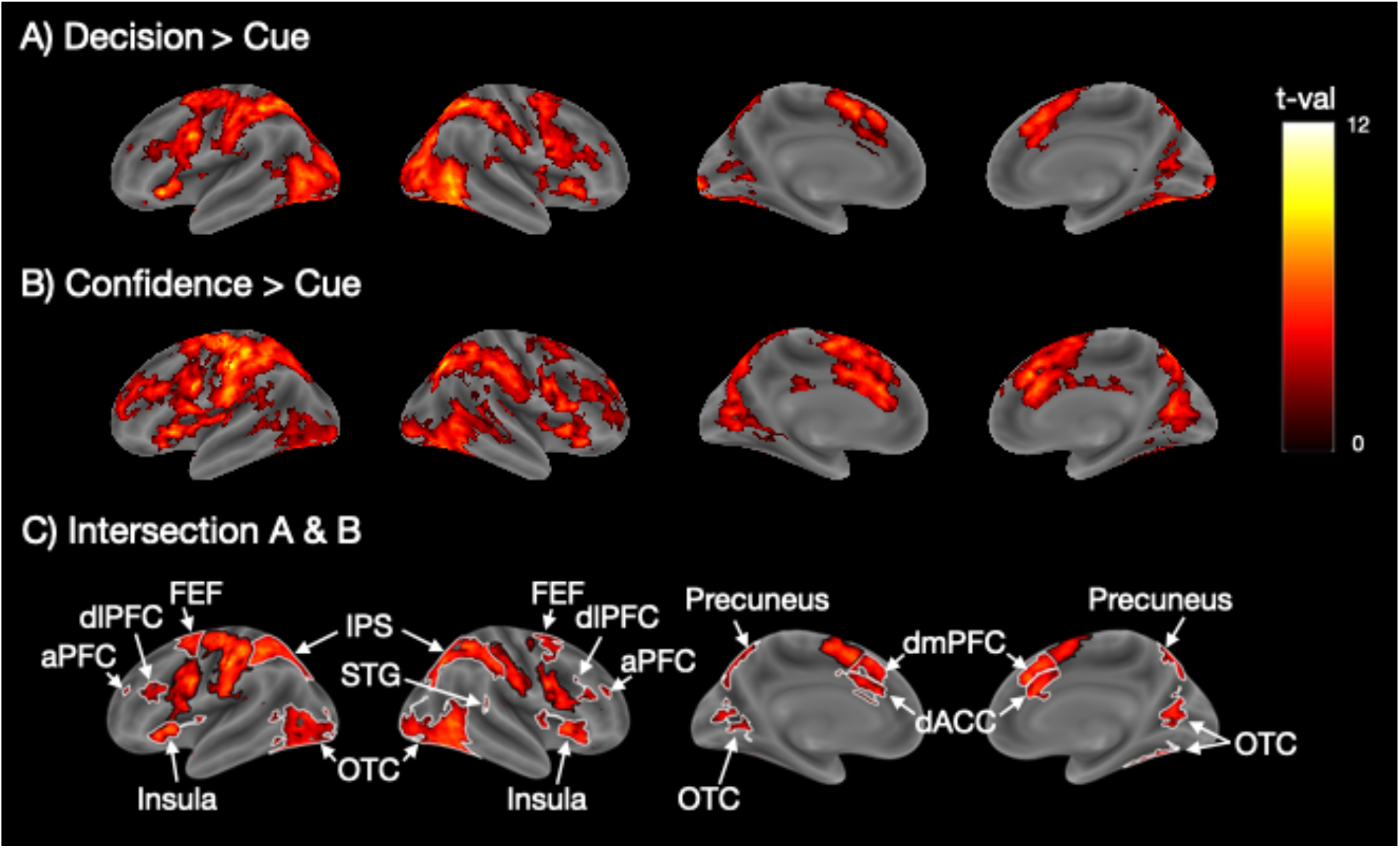
Shared activity between periods of perceptual decision making and confidence. (A) Decision-related brain activity obtained from the contrast Decision > Cue. (B) Confidence report-related brain activity obtained from the contrast Confidence > Cue. (C) Intersection between the activation maps for perceptual decision making and confidence showing the areas of activation overlap. Colors indicate t-values. The t-values in (C) are the average of the Decision > Cue and Confidence > Cue t-values. The black borders delineate motor-related activations in the somatosensory, motor, and pre-supplementary motor cortex, while the white borders delineate all other regions. aPFC, anterior prefrontal cortex; dACC, dorsal anterior cingulate cortex; dlPFC, dorsolateral prefrontal cortex; dmPFC, dorsomedial prefrontal cortex; FEF, frontal eye field; IPS, intraparietal sulcus; OTC, occipitotemporal cortex; STG, superior temporal gyrus.

These results suggest the presence of substantial overlap between the brain areas activated during periods of perceptual decision making and confidence report. However, an alternative explanation for this overlap is that the decision and confidence report periods may not have been perfectly separated in our GLM analyses and thus the overlap is a purely statistical confound, especially considering that the decision and confidence regressors showed a small but positive correlation (r = .16, SD = .105). Therefore, in a different set of analyses, we explored the activations produced by the decision period in trials where subjects did not provide a confidence rating (Decision_no_conf_). We then repeated all analyses with the Decision_no_conf_ regressor instead of the Decision regressor and still observed very similar results (Supplementary Figure 4).

We further performed a number of control analyses to ensure the robustness of our findings. First, we confirmed that our results were not due to the chosen baseline (i.e., the cue period) by running the same contrasts but using the fixation period (spanning the period from onset of the fixation cross until the onset of the cue) as the baseline. We observed similar activation patterns for both the decision and the confidence periods, as well as the intersection of the two sets of activations (Supplementary Figure 5). Second, we confirmed that our results were independent of the coherence level used and that similar results were obtained when only the low or only the high coherence levels were analyzed (Supplementary Figure 6). Third, we verified that our results did not depend on the confidence scale used and that the same results were obtained for each of the three different rating scales (Supplementary Figure 7).

#### Unique activations for decision and confidence periods

The analyses so far point to a considerable overlap between the brain regions activated by periods of perceptual decision making and confidence report. However, despite this overlap, it is possible that there are dissociable activations in a subset of brain regions. To check for the presence of such unique activations, we examined the set of brain regions activated more strongly for the decision period (using the contrast Decision > Confidence). We found activity in bilateral OTC (Figure 3A and Supplementary Table 2), which could be driven by the presence of moving-dot stimuli during the decision but not during the confidence period. We further examined the set of brain regions activated more strongly for the confidence period (using the contrast Confidence > Decision). We found activations in bilateral aPFC, dlPFC, dmPFC, dACC, IPL, precuneus, left insula, and a collection of regions near temporoparietal junction (TPJ) (Figure 3B and Supplementary Table 3). These results suggest the presence of an extensive network of regions that are more activated during the confidence period.

**Figure 3.**
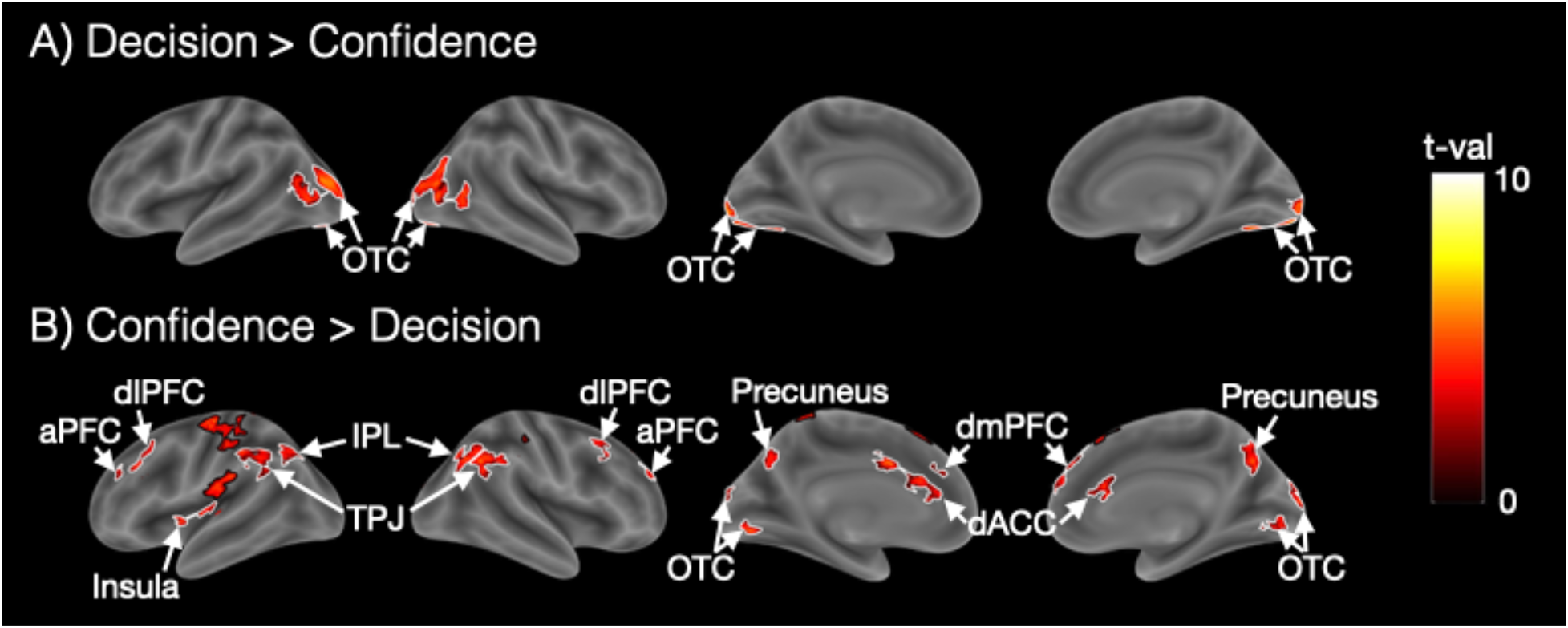
Unique activity for periods of perceptual decision making and confidence. (A) Regions showing higher activations for the decision than the confidence report period. (B) Regions showing higher activations for the confidence than the decision period. Colors indicate t-values.

The black borders delineate the somatosensory, motor, and pre-supplementary motor cortex, while the white borders delineate all other activations. aPFC, anterior prefrontal cortex; dACC, dorsal anterior cingulate cortex; dlPFC, dorsolateral prefrontal cortex; dmPFC, dorsomedial prefrontal cortex; FEF, frontal eye field; IPL, inferior parietal lobule; OTC, occipitotemporal cortex; TPJ, temporoparietal junction.

### Experiment 2

In Experiment 1, we separated the regressors associated with the periods of decision making and confidence report. However, despite the statistical separation of the regressors associated with each period, it is unlikely that Experiment 1 fully separated the actual processes of perceptual decision making and confidence generation. Therefore, to separate these processes better, we conducted Experiment 2 in which subjects completed the perceptual decisionmaking task without reporting confidence for the first half of the experiment, and only rated confidence in the second part of the experiment (Figure 1B). We call the first half of the experiment the “Confidence-absent” condition and the second half of the experiment the “Confidence-present” condition.

#### Behavioral results

The overall task accuracy was 74.2% (SD = .079) and the average response time was 728 ms (SD = 249). The average confidence across all trials was 2.866 (SD = .459) and average confidence RT was 516 ms (SD = 168). Confidence was higher for correct trials (2.953, SD = .454) than for incorrect trials (2.546, SD = .491; paired t-test: t(38) = 9.80, *p* = 6 x 10^-12^). These results suggest that the subjects performed the task well and evaluated their confidence level appropriately.

#### Shared activations between perceptual decision making and confidence generation

As in Experiment 1, we first examined the activity related to the decision and confidence periods separately. We examined the brain activity for the decision period from the Confidence-absent condition (Decision_conf-abs_), the decision period from the Confidence-present condition (Decision_conf-pres_), and the confidence period from the Confidence-present condition (Confidence).

We observed largely overlapping patterns of activation in fronto-parieto-posterior regions for all three of these periods (Figure 4A-C) even though the decisions in the Confidence-absent condition were made before subjects were ever told about rating confidence. Critically, as in Experiment 1, we created intersection maps between Decision_conf-abs_ and Confidence (Figure 4D and Supplementary Table 4), as well as between Decision_conf-pres_ and Confidence (Figure 4E and Supplementary Table 5). Both intersection maps were largely similar to what we observed in Experiment 1: there were shared activations in bilateral dACC, dlPFC, dmPFC, FEF, IPS, OTC, STG, insula, precuneus, and left aPFC. Therefore, the extensive overlap between the periods of perceptual decision making and confidence is both replicable for different experiments and can be found even when the decision period activity is estimated from trials that were completed before confidence ratings were ever introduced.

**Figure 4.**
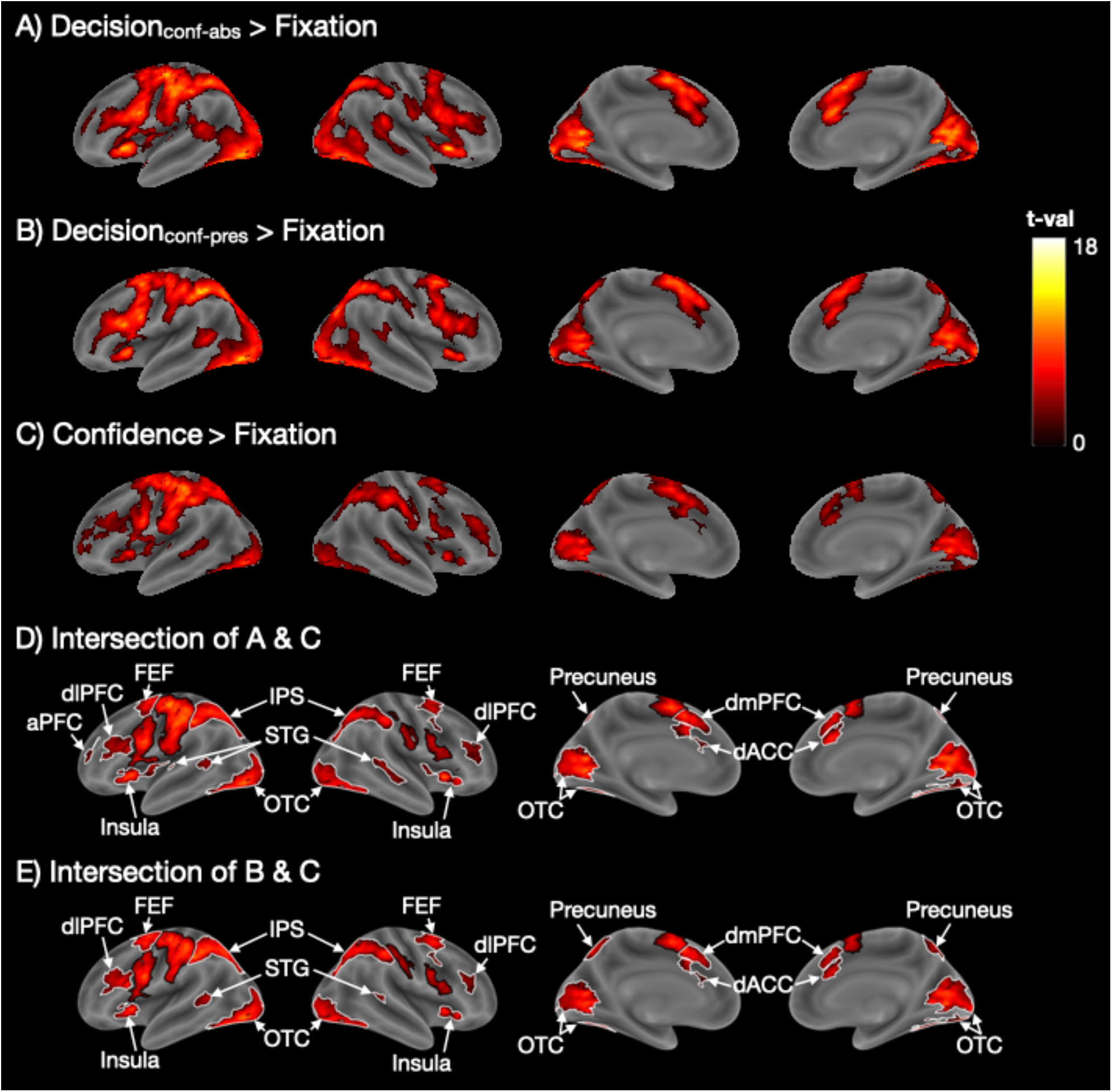
Overlapping brain activations for periods of perceptual decision making and confidence in Experiment 2. (A) Decision-related activity in the Confidence-absent condition (Decision_conf-abs_ > Fixation). (B) Decision-related activity in the Confidence-absent condition (Decision_conf-pres_ > Fixation). (C) Confidence-related activity (Confidence > Fixation). (D,E) Intersection maps for the decision-related and confidence-related activations. Colors indicate t-values. The t-values in (D) and (E) are the average t-values from the corresponding maps. The black borders delineate motor-related activations in the somatosensory, motor, and pre-supplementary motor cortex, while white borders delineate all other activations. aPFC, anterior prefrontal cortex; dACC, dorsolateral anterior cingulate cortex; dlPFC, dorsolateral prefrontal cortex; dmPFC, dorsomedial prefrontal cortex; FEF, frontal eye field; IPS, intraparietal sulcus; OTC, occipitotemporal cortex; STG, superior temporal gyrus.

#### Unique activations for perceptual decision making and confidence periods

As in Experiment 1, we examined the brain regions that were preferentially activated by either decision- or confidence-related periods. We first explored the regions that were activated more strongly during perceptual decision making by performing the contrasts Decision_conf-abs_ > Confidence and Decision_conf-pres_ > Confidence. As in Experiment 1, the strongest activation was in bilateral OTC (Figure 5A,B and Supplementary Table 6), which was likely caused by the moving dot stimuli. We also observed activations in medial prefrontal cortex (mPFC) and precuneus in both contrasts, as well as left STG and middle temporal gyrus (MTG) in the Decision_conf-abs_ > Confidence contrast though none of these regions appeared in the same contrast in Experiment 1.

**Figure 5.**
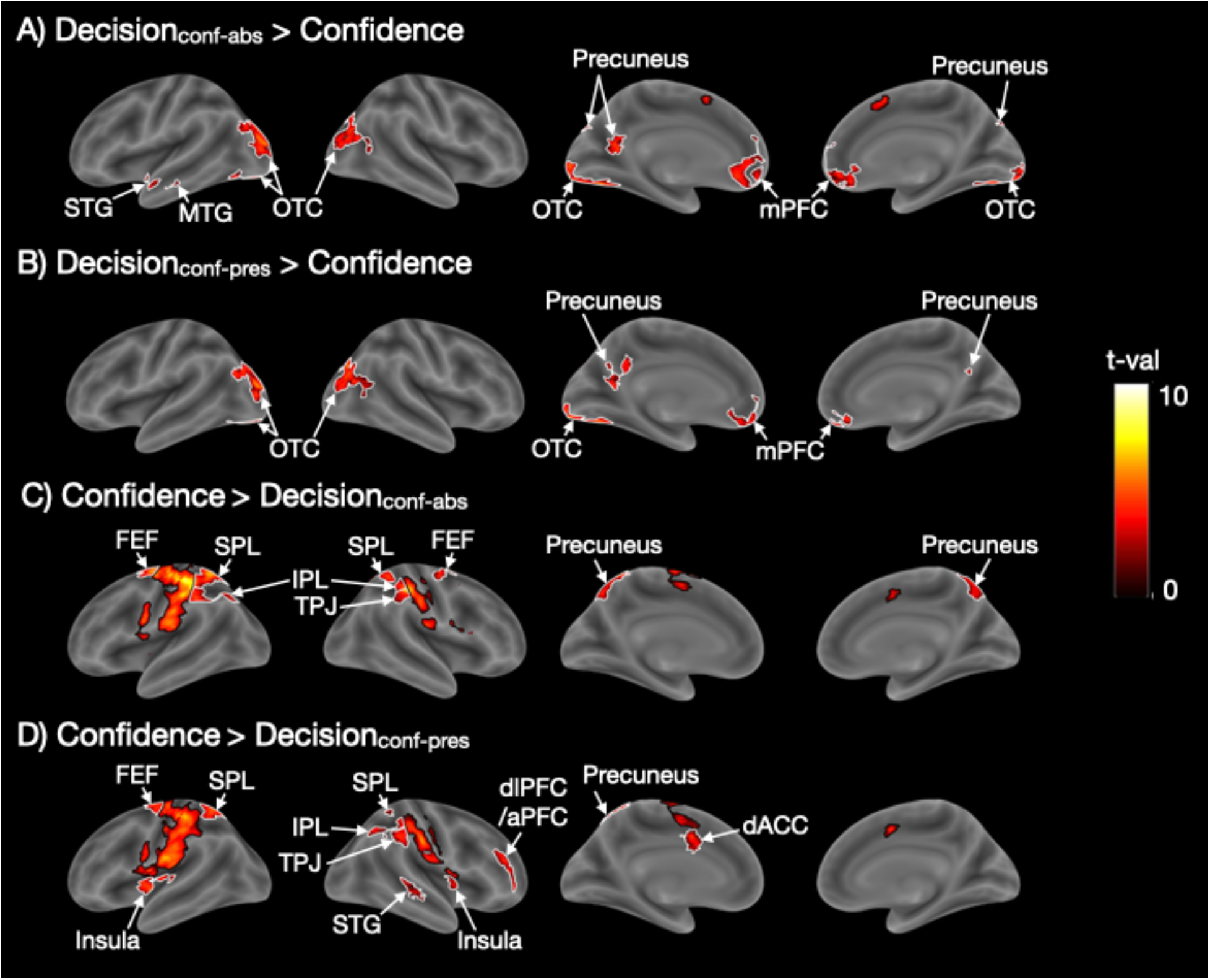
Unique brain regions activated for periods of perceptual decision making and confidence in Experiment 2. (A,B) Regions showing higher activations for Decision_conf-abs_ and Decision_conf-pres_ than Confidence. (C,D) Regions showing higher activations for Confidence than Decision_conf-abs_ and Decision_conf-pres_. Black borders delineate somatosensory, motor, and pre-supplementary motor cortex, while white borders delineate all other activations. Colors indicate t-values. aPFC, anterior prefrontal cortex; dACC, dorsal anterior cingulate cortex; dlPFC, dorsolateral prefrontal cortex; FEF, frontal eye field; IPL, inferior parietal lobule; mPFC, medial prefrontal cortex; MTG, middle temporal gyrus; OTC, occipitotemporal cortex; SPL, superior parietal lobule; STG, superior temporal gyrus; TPJ, temporoparietal junction.

Finally, as in Experiment 1, we examined what brain regions are more activated for confidence reporting compared to perceptual decision making. To do so, we contrasted the confidence report period with the decision period in both the Confidence-absent (Confidence > Decision_conf-abs_) and Confidence-present conditions (Confidence > Decision_conf-pres_). The contrast Confidence > Decision_conf-abs_ (Figure 5C and Supplementary Table 7) revealed activations in bilateral IPL, SPL, FEF, precuneus, and right TPJ. While most of those areas activated in Confidence > Decision_conf-pres_ (Figure 5D and Supplementary Table 8), we found some extra brain regions like bilateral insula, right dlPFC/aPFC, right STG, and left dACC. In addition, we observed activations in motor and somatosensory cortex that are likely related to the button presses and not the confidence reporting process itself. In control analyses, we performed the same contrasts but not only for the Neutral blocks (where neutral cues were given) but for all trials (including blocks with predictive cues). This analysis tripled the number of trials and thus provided more power. Consequently, we observed more extensive activations, including extensive activations in bilateral aPFC/dlPFC and anterior insula (Supplementary Figure 8). Overall, there were a number of similarities between the areas that were more activated for confidence report in Experiments 1 and 2.

## Discussion

We investigated whether the periods of perceptual decision making and confidence activate the same or different brain regions. In two experiments, we found extensive overlap between the brain circuits activated during the decision and confidence report periods. At the same time, a network of frontal, temporo-parietal, and cingulate regions was more active during the confidence period, but the only OTC was more active for the decision than the confidence period across the two experiments. These findings suggest that perceptual decision making and confidence reports are generated by largely overlapping brain circuits though the confidence period recruits additional areas.

### What is the reason for the activation overlap?

We found extensive overlap between regions that were active during the decision and confidence periods in both of our experiments. Shared activations emerged in frontal (aPFC, dlPFC, dmPFC, FEF, and insula), parietal (IPL, and STG), posterior (OTC and precuneus), and cingulate (dACC) regions. A critical question, therefore, is what caused the consistent activation overlap. There appear to be two broad, non-mutually-exclusive possibilities that we explore below. While we favor the first option, it is possible that the second option also contributes to our results.

#### Option 1: Perceptual decision making and confidence generation genuinely activate similar brain areas

One possible interpretation of our findings is that the process behind perceptual decision making and confidence genuinely activate a very similar set of brain areas. This possibility can be seen as unsurprising; after all, both processes are effortful^62^ and involve the transformation of an internal signal into a judgment (perceptual or metacognitive) that is eventually expressed using a button press. Indeed, these similarities make some degree of overlap virtually inevitable. Nevertheless, this is the first study to show just how thorough the overlap between the activations related to the periods of perceptual decision making and confidence really is.

Overlap between perceptual decision making and confidence has been demonstrated for a number individual regions in animal studies^7,13–16^. These studies have shown that the same populations of neurons predict both the decision and the confidence in the decision. Nevertheless, these previous studies did not attempt to separate the decision and confidence computations temporally. For example, in the common opt-out task, animals are allowed to choose a safe option that results in certain but small reward^63^. This common paradigm features a complete temporal overlap between perceptual decision making and confidence generation, making it challenging to ensure that neural activity is specifically related to decision or confidence.

The possibility that perceptual decision making and confidence generation genuinely activate similar brain areas would imply that perceptual decisions and confidence judgments may not be as fundamentally different from each other as sometimes assumed. For example, it is common to refer to these judgments as Type 1 vs. Type 2 because the perceptual decision is about the stimulus (Type 1 judgment), whereas the confidence judgment is about the accuracy of the decision (Type 2 judgment)^64^. This terminology may suggest that confidence judgments are completely different than perceptual decision making. However, another view that comes from the early days of signal detection theory is that confidence judgments can be conceptualized as simply another perceptual decision that uses different decision criteria on the same underlying signal^65^. This conceptualization implies that perceptual decision making and confidence are in fact quite similar. Although the current brain results address this question only indirectly, they can be seen as implying that the processes related to perceptual decision making and confidence generation are indeed more similar than different from each other.

Nevertheless, it is important to note that even if perceptual decision making and confidence generation were to activate the exact same brain areas, the actual computations that underlie these two processes may still be very different. Such differences are implied by accuracy-confidence dissociations^21,22,27–35^, neurological phenomena such as in blindsight^66^, and studies that delivered transcranial magnetic stimulation (TMS) to the early visual cortex and found separable effects on accuracy and confidence^67–69^. Moreover, a recent study provided direct evidence that sensory neural signals are indeed used in different ways in making perceptual decisions and confidence judgments^24^. The confidence-accuracy dissociations in previous studies imply the existence of at least partially separate neural computations for perceptual decision making and confidence.

#### Option 2: Decision- and confidence-related activity take place at the same time

A second, non-exclusive possibility is that the extensive overlap between the activations related to the decision and confidence periods is due to the decision- and confidence-related processes having a substantial temporal overlap. According to this possibility, not only were decision- and confidence-related processes not confined to the decision and confidence periods, but at least one process (and potentially both) was substantially engaged during both periods.

Our experiments were designed to exclude the possibility of trivial findings where the overlap is driven by lack of separation of the fMRI regressors themselves. For example, in Experiment 1, we separated the fMRI regressors for the decision and confidence report periods. Experiment 2 had an even stronger manipulation where confidence was never introduced in the first half of the experiment. It is clear that these manipulations were at least partially successful. First, we observed large bilateral occipital activations in the decision period compared to the confidence period in both experiments. The activation would be expected given that the moving dot stimulus was present during the decision but not the confidence period. Second, the regions activated preferentially for the confidence period are known to be part of a network of regions involved in confidence^2,40,41,70–75^. Neither of these results would be expected if we did not achieve at least partial separation between the periods of perceptual decision making and confidence report. Nevertheless, these results do not exclude the possibility that some confidence report processes may have taken place during the decision period or that some decision processes may have taken place during the confidence period (or both).

Overall, our results do not allow us to isolate the exact reasons for the strong overlap between decision and confidence period activations and thus further research is needed to convincingly settle this issue. Nevertheless, our results point in the direction that the periods of perceptual decision making and confidence report are likely to genuinely activate a very similar set of brain areas.

### Brain areas more active in confidence judgments

Despite the extensive overlap of the activations for the decision and the confidence periods, we also found a number of brain regions that were more active during the confidence compared to the decision period. These regions appeared in both experiments and were located in the prefrontal cortex (aPFC, dlPFC, and dmPFC), the dorsal anterior cingulate cortex (dACC), precuneus, insula, and near the temporoparietal junction (TPJ and IPL). Many of these areas have been linked to confidence computations in previous studies^2,40,41,70–75^ but it has remained unclear whether such activations are stronger than activations that may be caused by the perceptual decision itself. For example, dlPFC has been implicated in both the perceptual decision^76–79^ and confidence^27,41,45,80^. Our results suggest that at least a subregion of dlPFC is in fact more active during the period of confidence report compared to the decision period.

Why are some regions more active during the confidence period? One possibility is that judgments of confidence serve not only as a guide to the external environment but also reflect our own internal states. For example, repeatedly having low confidence may indicate not only that the stimulus is difficult but that one is losing alertness. Such process of self-evaluation could be supported by areas in and around TPJ that are known to be involved in processing social information^81,82^. Further, confidence evaluations are used to alter one’s strategy for doing the task on subsequent trials^22^ and are thus linked to control processes in the frontal and cingulate regions.

### Are there any regions more active for perceptual decision making than confidence report?

The only region, that was more active during the decision period than the confidence period across the two experiments was bilateral occipitotemporal cortex (OTC). Activity in OTC is likely to be driven by the moving dot stimuli and not by the decision-making process itself (though it is impossible to completely exclude this possibility without further experiments where the sensory stimulation remains the same during the periods of decision making and confidence report). We also observed several activations that were specific to just one experiment such as bilateral mPFC and precuneus in Experiment 2. These two specific regions have previously been found to associated with subjective confidence^74,83^ and it is thus surprising that they were activated preferentially for the decision period in Experiment 2. Nevertheless, none of these regions was activated in both Experiments 1 and 2, and thus they are unlikely to be preferentially involved in perceptual decision making independent of the specifics of each experimental paradigm.

The lack of regions that are clearly selective for the decision itself could be interpreted as suggesting that perceptual decision making does not involve unique process components not present in confidence judgments. Instead, it could be argued that the processes unfolding during the confidence report completely subsume the processes unfolding during the decision period (and include additional components). Further research is needed in order to support (or falsify) this possibility.

### Conclusion

We found a surprisingly high degree of overlap in the brain regions activated during periods of perceptual decision making and confidence report. In addition, unique brain activations were found for the confidence period but not for the perceptual decision-making period. The inclusion of two experiments with different designs also helped establish which activations are robust across non-trivial changes in the experimental paradigm and which activations appear to be specific to the experimental design used in each experiment. We suggest that future research should adopt a similar approach where the overlap between the brain areas supporting various processes should be examined across several different contexts.

## Author contributions

All authors designed the studies. J. Yeon and M. Shekhar collected the data. J. Yeon analyzed the data, and J. Yeon and D. Rahnev worked together to interpret the results. J. Yeon drafted the manuscript; D. Rahnev and M. Shekhar revised and commented on it. All authors approved the final version of the manuscript for submission. The authors declare no competing interests.

## Acknowledgements

We thank Farshad Rafiei for help with the data collection. This work was supported by the National Institutes of Health (Award Number: R01MH119189) and the Office of Naval Research (Award Number: N00014-20-1-2622).

