## Supplementary materials for "Overlapping and unique neural circuits are activated during perceptual decision making and confidence"

### Supplementary Figures

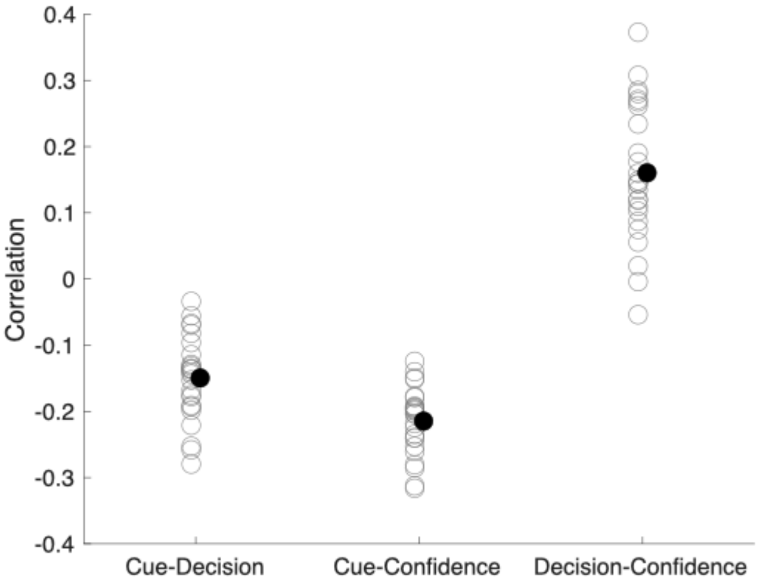


**Supplementary Figure 1.** Correlation values between different regressors in Experiment 1. Open circles indicate individual subjects' data and the closed black circle shows the average across the subjects.

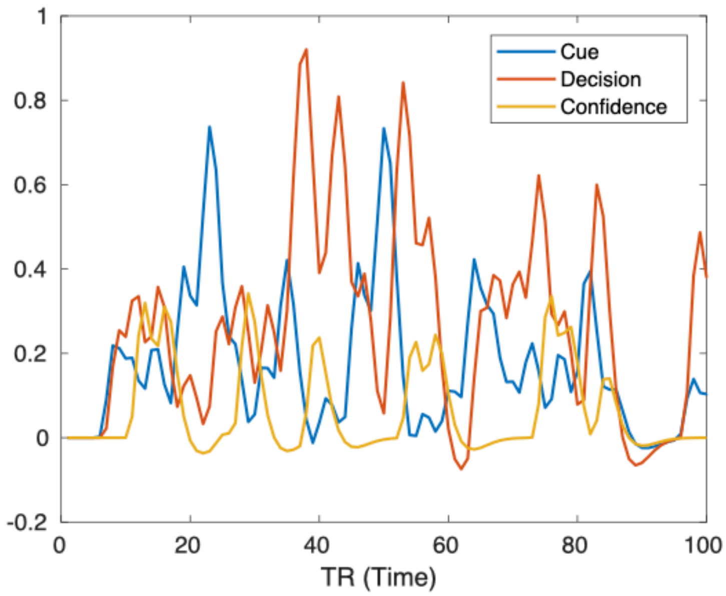


**Supplementary Figure 2.** Time-series for the regressors for the Cue, Decision, and Confidence periods from a single subject. As can be appreciated visually, the time courses for the regressors are fairly distinct.

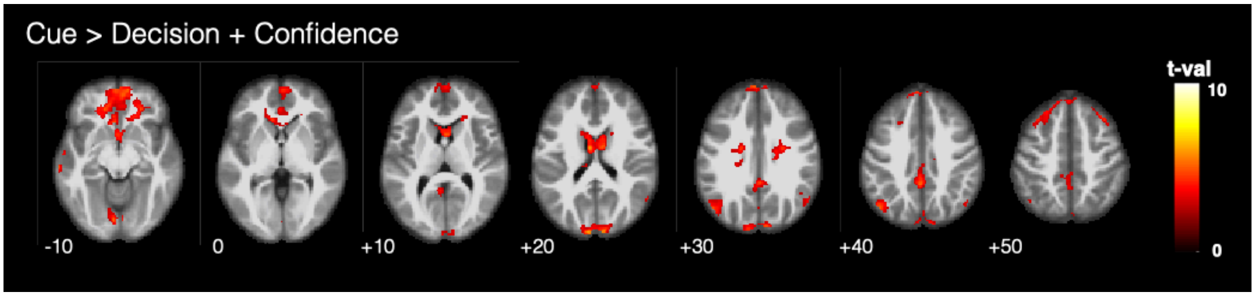


**Supplementary Figure 3**. Cue-related activity. The brain activations during the cue period were assessed using the contrast Cue > Decision + Confidence. The cue activated the default mode network but not known attention-related regions such as the dorsal attention network. Activation patterns displayed using a $p < .001$ uncorrected threshold. Only clusters larger than 5 voxels are displayed. The colors indicate t-values.

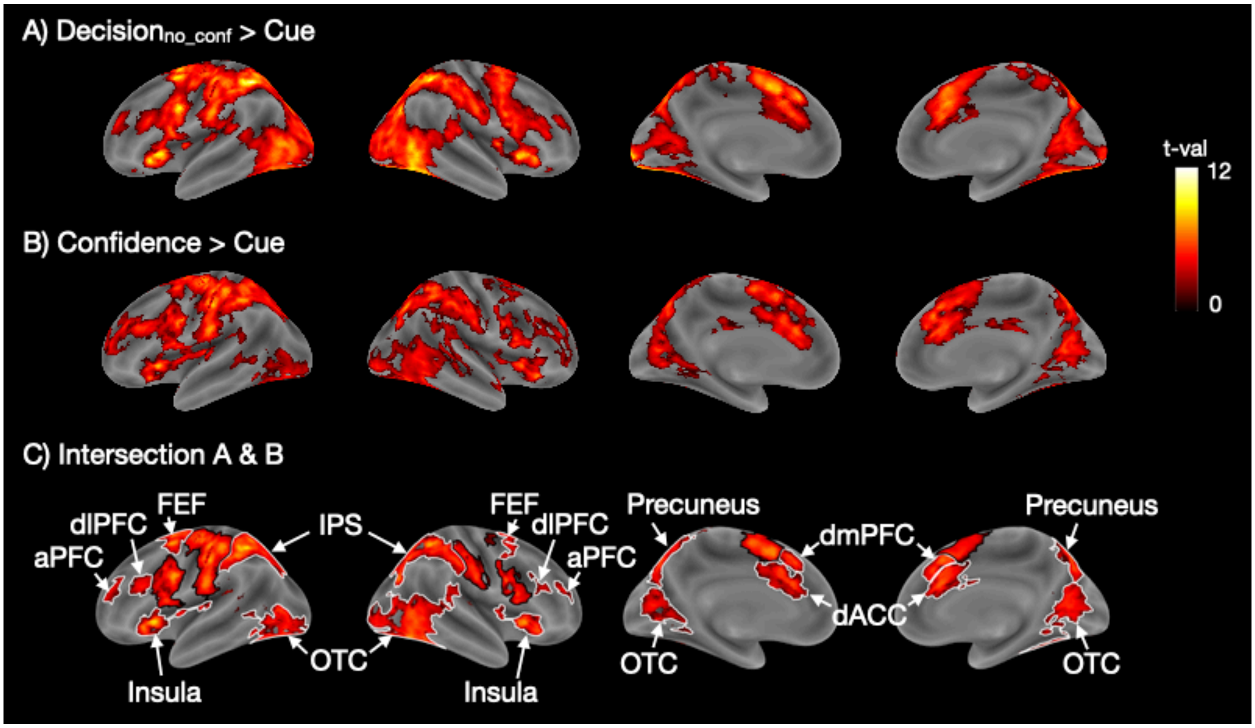


**Supplementary Figure 4**. Shared activity between periods of decision and confidence where the decision period activity was estimated based only on trials that were not followed by a confidence report. (A) Decision-related brain activity obtained from the contrast Decision$_{no_conf}$ > Cue. (B) Confidence report-related brain activity obtained from the contrast Confidence > Cue. (C) Intersection between the activation maps for perceptual decision making and confidence showing the areas of activation overlap. Overall, the results indicate very similar pattern of results to our main analyses (Figure 2) where the decision-related activity was estimated from all trials. Black borders delineate somatosensory, motor, and pre-supplementary motor cortex, while white borders delineate all other activations. Colors indicate t-values. The t-values in (C) are the average of the Decision$_{no_conf}$ > Cue and Confidence > Cue t-values. aPFC, anterior prefrontal cortex; dACC, dorsal anterior cingulate cortex; dlPFC, dorsolateral prefrontal cortex; FEF, frontal eye field; IPS, inferior parietal sulcus; OTC, occipitotemporal cortex.

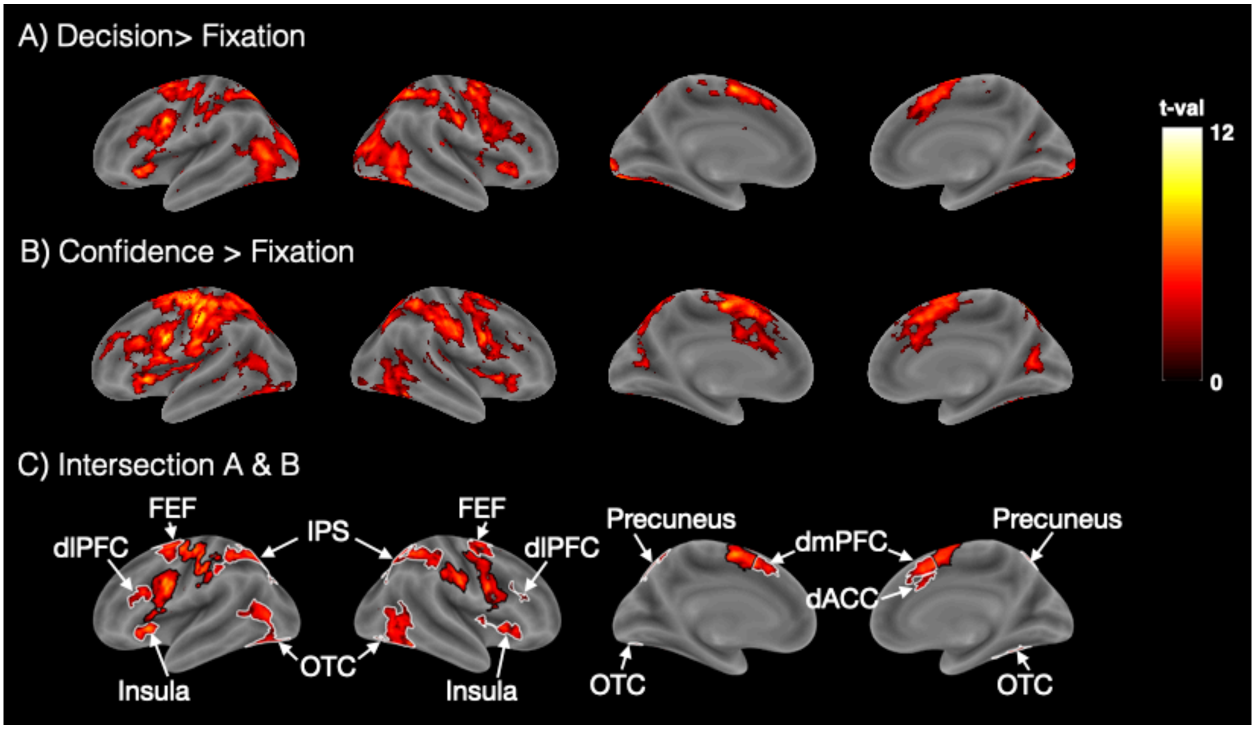


**Supplementary Figure 5**. Control analyses in which the decision- and confidence report-related activity was assessed by using the fixation – rather than the cue – period as a baseline. (A) Decision-related brain activity obtained from the contrast Decision > Fixation. (B) Confidence report-related brain activity obtained from the contrast Confidence > Fixation. (C) Intersection between the activation maps for perceptual decision making and confidence showing the areas of activation overlap. All of the results above are similar to the results obtained when the cue period was used as a baseline (Figure 2). The colors indicate t-values. The t-values in (C) are the average of the Decision > Cue and Confidence > Cue t-values. The black borders delineate the somatosensory and motor cortices, and pre-supplementary motor area. dACC, dorsal anterior cingulate cortex; dlPFC, dorsolateral prefrontal cortex; dmPFC, dorsomedial prefrontal cortex; FEF, frontal eye field; IPS, Intraparietal sulcus; OTC, Occipitotemporal cortex.

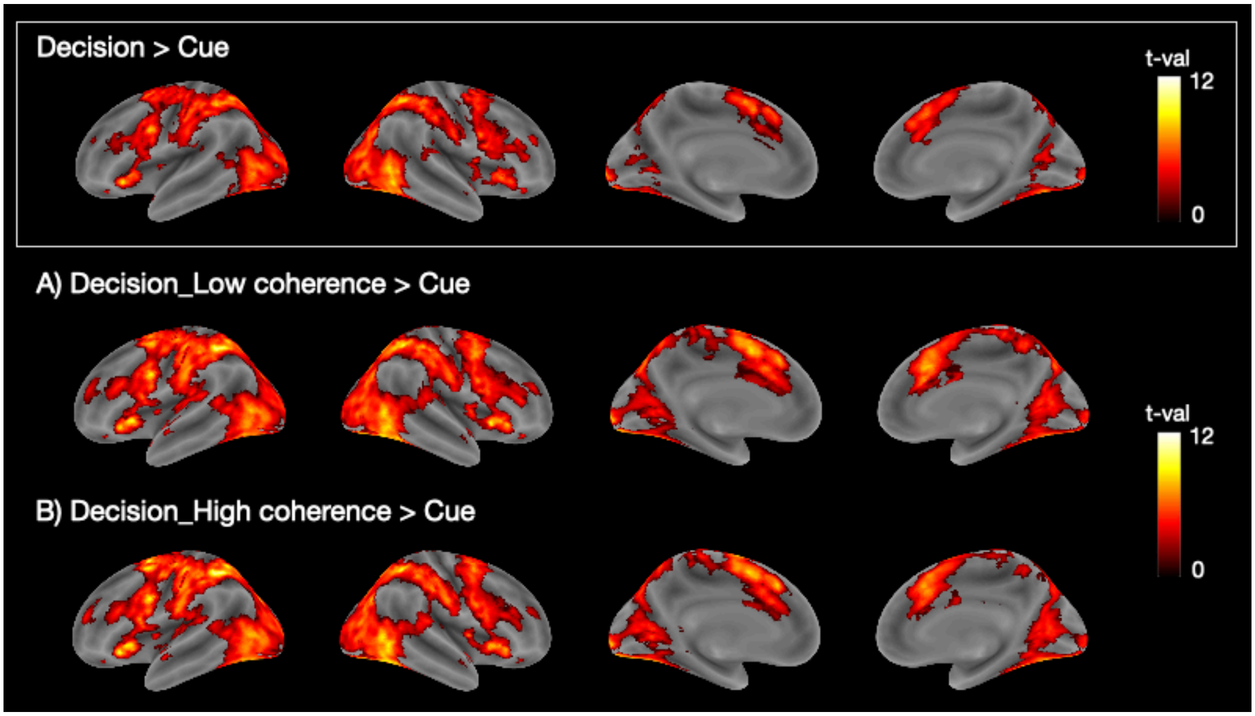


**Supplementary Figure 6**. Decision-related brain activity for (A) low-coherence, and (B) high-coherence stimuli. Top row (white box) shows Decision > Cue contrast result from the main analysis (i.e., decision-related brain activity without separating the coherence levels) for comparison. The activation patterns for the low- and high-coherence stimuli are similar to the activation pattern from the main analysis. The colors indicate t-values.

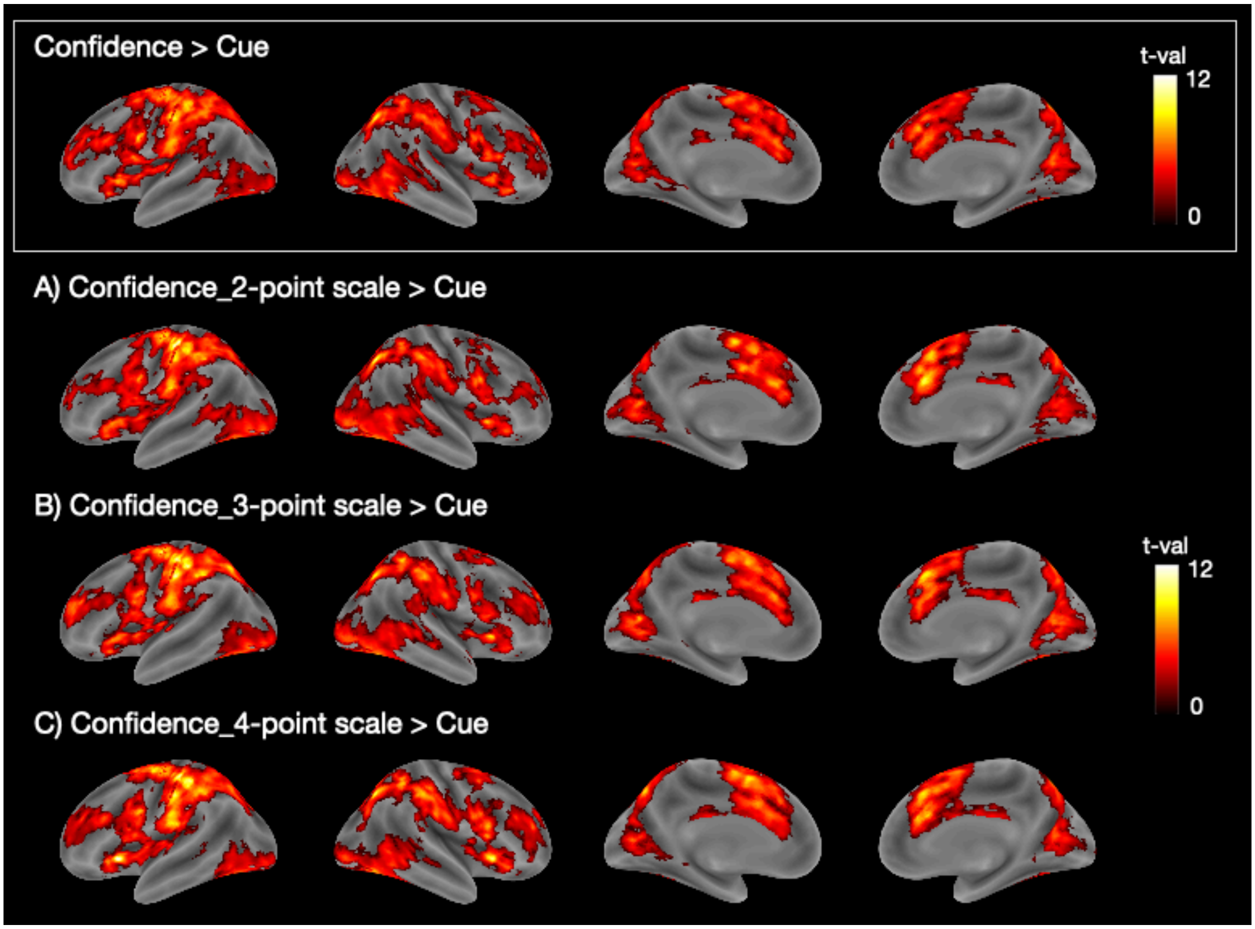


**Supplementary Figure 7**. Confidence-related brain activity for the three confidence scales. Activated brain regions for the (A) 2-point, (B) 3-point, and (C) 4-point scales. Top row (white box) shows Confidence > Cue contrast result from the main analysis (i.e., confidence-related brain activity without separating the confidence scales) for comparison. The three activation patterns from the different rating scales are similar to the activation pattern from the main analysis. The colors indicate t-values.

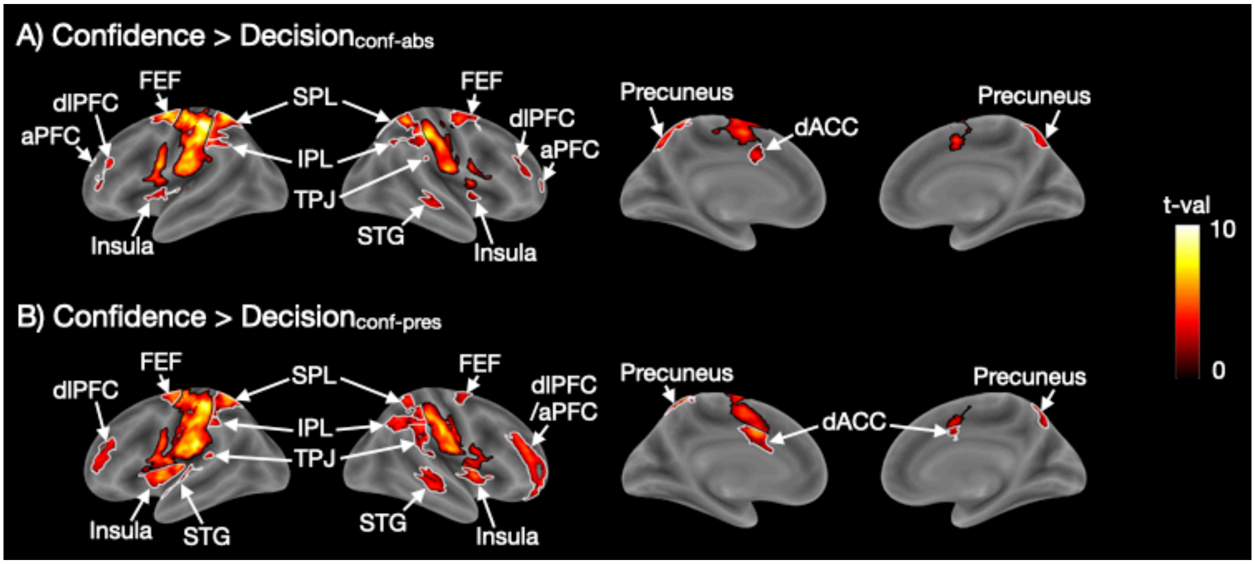


**Supplementary Figure 8.** Brain regions preferentially activated during the confidence period in Experiment 2 when all trials are considered. The main analyses in Experiment 2 focused only on a subset of trials where Neutral predictive cues were given. In this control analysis, we examined all trials, including ones where the cue predicted either coherent or random motion. We found similar results as in the main analyses (see Figure 5) but the greater power produced larger areas of activation. Black borders delineate somatosensory, motor, and pre-supplementary motor cortex, while white borders delineate all other activations. Colors indicate t-values. aPFC, anterior prefrontal cortex; dACC, dorsal anterior cingulate cortex; dlPFC, dorsolateral prefrontal cortex; FEF, frontal eye field; IPL, inferior parietal lobule; SPL, superior parietal lobule; STG, superior temporal gyrus; TPJ, temporoparietal junction.

### Supplementary Tables

| | | MNI Coordinates | | | t-value | |
|---|---|---|---|---|---|---|
| Anatomical region | Side | x | y | z | Decision > Cue | Confidence > Cue |
| Dorsomedial prefrontal cortex (dmPFC) | L | -4 | 12 | 50 | 9.36 | 6.88 |
| | R | 4 | 22 | 40 | 6.80 | 9.34 |
| Frontal eye field (FEF) | L | -26 | -2 | 56 | 9.40 | 6.99 |
| | R | 30 | -2 | 56 | 8.02 | 4.69 |
| Intraparietal sulcus (IPS) | L | -40 | -32 | 42 | 6.98 | 9.33 |
| | R | 38 | -40 | 46 | 7.87 | 9.20 |
| Occipito temporal cortex (OTC) | L | -50 | -76 | 4 | 8.33 | 3.74 |
| | R | 54 | -68 | -2 | 9.71 | 5.30 |
| Insula | L | -30 | 24 | 2 | 6.58 | 7.56 |
| | R | 32 | 20 | 6 | 7.17 | 6.64 |
| Dorsal anterior cingulate cortex (dACC) | L | -6 | 24 | 32 | 4.54 | 7.99 |
| | R | 10 | 28 | 28 | 5.61 | 8.46 |
| Precuneus | L | -14 | -74 | 56 | 7.21 | 4.82 |
| | R | 14 | -68 | 56 | 5.56 | 7.51 |
| Dorsolateral prefrontal cortex (dlPFC) | L | -46 | 22 | 24 | 3.91 | 5.10 |
| | R | 52 | 28 | 22 | 4.02 | 5.63 |
| Anterior prefrontal cortex (aPFC) | L | -30 | 44 | 22 | 3.07 | 3.88 |
| | R | 34 | 44 | 22 | 3.55 | 4.39 |
| Superior temporal gyrus (STG) | R | 60 | -38 | 14 | 3.33 | 3.04 |

**Supplementary Table 1**. Coordinates and t-values for the peak voxel of each activated cluster for the intersection of the Decision > Cue and the Confidence > Cue contrasts in Experiment 1.

| Anatomical region | Side | MNI Coordinates x | y | z | t-value |
|---|---|---|---|---|---|
| Occipitotemporal cortex (OTC) | L | -44 | -88 | 14 | 6.57 |
| | R | 30 | -94 | 12 | 6.41 |

**Supplementary Table 2**. Coordinates and t-values for the peak voxel of each activated cluster for the Decision > Confidence contrast in Experiment 1.

| | | MNI Coordinates | | | |
|---|---|---|---|---|---|
| Anatomical region | Side | x | y | z | t-value |
| Insula | L | 22 | 48 | 32 | 6.86 |
| | R | -30 | 42 | 34 | 6.56 |
| Dorsal anterior cingulate cortex (dACC) | L | -2 | 8 | 36 | 6.50 |
| | R | 4 | 26 | 18 | 5.07 |
| Occipitotemporal cortex (OTC) | L | -10 | -80 | -4 | 5.38 |
| | R | 8 | -70 | -8 | 6.31 |
| Dorsolateral prefrontal cortex (dlPFC) | L | -36 | 24 | 30 | 6.18 |
| | R | 36 | 20 | 42 | 4.78 |
| Temporoparietal junction (TPJ) | L | -50 | -48 | 52 | 5.36 |
| | R | 50 | -54 | 42 | 6.03 |
| Inferior parietal lobule (IPL) | L | -26 | -44 | 68 | 5.67 |
| | R | 48 | -42 | 54 | 4.73 |
| Precuneus | L | -2 | -74 | 42 | 4.79 |
| | R | 4 | -70 | 36 | 5.18 |
| Dorsomedial prefrontal cortex (dmPFC) | L | -2 | 32 | 38 | 4.56 |
| | R | 10 | 40 | 44 | 4.50 |
| Anterior prefrontal cortex (aPFC) | L | -30 | 34 | 24 | 3.97 |
| | R | 12 | 52 | 22 | 4.65 |

**Supplementary Table 3**. Coordinates and t-values for the peak voxel of each activated cluster for the Confidence > Decision contrast in Experiment 1.

| Anatomical region | Side | MNI Coordinates | | | t-value | |
|---|---|---|---|---|---|---|
| | | | | | Confidence-absent | Confidence-present |
| | | x | y | z | Decision > Fixation | Confidence > Fixation |
| Insula | L | -34 | 14 | 2 | 10.48 | 7.22 |
| | R | 34 | 20 | 0 | 13.54 | 5.89 |
| Dorsal anterior cingulate cortex (dACC) | L | -8 | 14 | 36 | 9.25 | 8.93 |
| | R | 8 | 22 | 30 | 10.88 | 4.44 |
| Intraparietal sulcus (IPS) | L | -48 | -36 | 46 | 10.61 | 9.88 |
| | R | 44 | -34 | 42 | 7.41 | 7.82 |
| Frontal eye field (FEF) | L | -26 | -4 | 54 | 10.75 | 8.12 |
| | R | 32 | 0 | 58 | 5.78 | 7.51 |
| Dorsolateral prefrontal cortex (dlPFC) | L | -46 | 30 | 24 | 4.48 | 4.75 |
| | R | 52 | 36 | 24 | 3.57 | 5.59 |
| Superior temporal gyrus (STG) | L | -52 | -50 | 8 | 5.82 | 3.52 |
| | R | 46 | -28 | -4 | 4.41 | 6.50 |
| Anterior prefrontal cortex (aPFC) | L | -34 | 54 | 20 | 3.16 | 3.55 |
| Occipito temporal cortex (OTC) | L | -42 | -84 | -14 | 11.62 | 9.00 |
| | R | 36 | -84 | -14 | 7.63 | 5.07 |
| Precuneus | L | -12 | -68 | 48 | 3.59 | 7.04 |
| | R | 16 | -68 | 52 | 5.74 | 7.10 |
| Dorsomedial prefrontal cortex (dmPFC) | L | -6 | 12 | 48 | 12. 12 | 7.20 |
| | R | 6 | 12 | 50 | 16.19 | 3.81 |

**Supplementary Table 4**. Coordinates and t-values for the peak voxel of each activated cluster for the intersection of the $\text{Decision}_{\text{conf-abs}}$ > Fixation and Confidence > Fixation contrasts in Experiment 2.

| Anatomical region | Side | MNI Coordinates | | | t-value Confidence-present | |
|---|---|---|---|---|---|---|
| | | x | y | z | Decision > Fixation | Confidence > Fixation |
| Dorsomedial prefrontal cortex (dmPFC) | L | -2 | 14 | 44 | 15.41 | 8.20 |
| | R | 2 | 14 | 48 | 13.34 | 6.26 |
| Occipito temporal cortex (OTC) | L | -42 | -82 | -12 | 12.78 | 8.12 |
| | R | 40 | -86 | -16 | 8.93 | 5.34 |
| Frontal eye field (FEF) | L | -24 | -4 | 54 | 12.34 | 8.54 |
| | R | 32 | -2 | 56 | 9.33 | 6.69 |
| Dorsal anterior cingulate cortex (dACC) | L | -8 | 12 | 46 | 11.68 | 7.05 |
| | R | 8 | 26 | 32 | 6.80 | 6.90 |
| Intraparietal sulcus (IPS) | L | -48 | -36 | 44 | 11.28 | 9.67 |
| | R | 42 | -34 | 40 | 8.90 | 8.35 |
| Insula | L | -34 | 14 | 2 | 10.25 | 7.22 |
| | R | 34 | 20 | 0 | 9.24 | 5.89 |
| Dorsolateral prefrontal cortex (dlPFC) | L | -52 | 20 | 26 | 7.24 | 3.55 |
| | R | 52 | 12 | 28 | 8.60 | 5.45 |
| Precuneus | L | -16 | -68 | 56 | 8.11 | 7.04 |
| | R | 16 | -68 | 54 | 6.48 | 7.58 |
| Superior temporal gyrus (STG) | L | -48 | -50 | 4 | 7.24 | 3.03 |
| | R | 50 | -38 | 8 | 3.93 | 4.43 |

**Supplementary Table 5**. Coordinates and t-values for the peak voxel of each activated cluster for the intersection of the $\text{Decision}_{\text{conf-pres}}$ > Fixation and Confidence > Fixation contrasts in Experiment 2.

**Decision_conf-abs > Confidence**

| Anatomical region | Side | MNI Coordinates x | y | z | t-value |
|---|---|---|---|---|---|
| Occipito temporal cortex (OTC) | L | -24 | -94 | 14 | 5.98 |
| | R | 38 | -84 | 26 | 5.59 |
| Precuneus | L | -10 | -54 | 10 | 6.59 |
| | R | 22 | -80 | 48 | 3.80 |
| Middle prefrontal cortex (mPFC) | L | -2 | 34 | -14 | 5.31 |
| | R | 2 | 46 | -2 | 5.04 |
| Middle temporal gyrus (MTG) | L | -62 | -6 | -18 | 6.14 |
| Superior temporal gyrus (STG) | L | -34 | 10 | -22 | 5.23 |

**Decision_conf-pres > Confidence**

| Anatomical region | Side | MNI Coordinates x | y | z | t-value |
|---|---|---|---|---|---|
| Occipito temporal cortex (OTC) | L | -28 | -90 | 16 | 6.27 |
| | R | 30 | -70 | 22 | 6.77 |
| Precuneus | L | -8 | -54 | 10 | 5.64 |
| | R | 4 | -60 | 20 | 3.86 |
| Middle prefrontal cortex (mPFC) | L | -2 | 52 | -4 | 5.19 |
| | R | 2 | 50 | -4 | 4.80 |

**Supplementary Table 6**. Coordinates and t-values for the peak voxel of each activated cluster for the $Decision_{conf-abs}$ > Confidence (top) and $Decision_{conf-pres}$ > Confidence (bottom) contrasts in Experiment 2.

|  |  | MNI Coordinates |  |  |  |
| --- | --- | --- | --- | --- | --- |
| Anatomical region | Side | x | y | z | t-value |
| Frontal eye field (FEF) | L | -26 | -4 | 64 | 6.60 |
|  | R | 30 | 2 | 60 | 4.44 |
| Superior parietal lobule (SPL) | L | -28 | -54 | 66 | 6.63 |
|  | R | 28 | -50 | 66 | 4.26 |
| Inferior parietal lobule (IPL) | L | -50 | -36 | 48 | 6.09 |
|  | R | 46 | -36 | 52 | 4.28 |
| Temporo-parietal junction (TPJ) | R | 52 | -36 | 46 | 3.22 |
| Precuneus | L | -10 | -72 | 50 | 4.40 |
|  | R | 10 | -68 | 50 | 3.54 |

**Supplementary Table 7**. Coordinates and t-values for the peak voxel of each activated cluster for the Confidence > Decision$_{conf\text{-}abs}$ contrast in Experiment 2.

| Anatomical region | Side | MNI Coordinates x | y | z | t-value |
|---|---|---|---|---|---|
| Superior parietal lobule (SPL) | L | -18 | -54 | 64 | 5.74 |
| | R | 16 | -68 | 58 | 3.88 |
| Insula | L | -44 | -6 | 4 | 5.64 |
| | R | 44 | -2 | 10 | 5.34 |
| Inferior parietal lobule (IPL) | R | 46 | -28 | 34 | 5.50 |
| Temporo-parietal junction (TPJ) | R | 52 | -40 | 42 | 4.47 |
| Frontal eye field (FEF) | L | -24 | -8 | 70 | 5.43 |
| Dorsal anterior cingulate cortex (dACC) | L | -4 | -2 | 48 | 5.39 |
| Dorsolateral/anterior prefrontal cortex (dlPFC/aPFC) | R | 40 | 48 | 24 | 5.23 |
| Precuneus | L | -16 | -58 | 64 | 4.75 |
| Superior temporal gyrus (STG) | R | 46 | -28 | -4 | 4.23 |

**Supplementary Table 8**. Coordinates and t-values for the peak voxel of each activated cluster for the Confidence > Decision$_{conf\text{-}pres}$ contrast in Experiment 2.
